# Antennal Lobe Dynamics And The Generation Of Diverse Response Patterns To Mechanosensory Stimulation

**DOI:** 10.64898/2026.09.09.750382

**Authors:** Joseph Reed, Mainak Patel

**Affiliations:** Department of Mathematics, William & Mary; Williamsburg, VA, 23185; USA

**Keywords:** olfaction, sensory integration, antennal lobe model, SK channel, slow inhibition, mechanosensory

## Abstract

Sensory integration within antennal lobe (AL) is thought to play a role in the guidance of odor tracking, though while responses to olfactory input within the AL have been well-studied, responses to mechanosensory stimuli, in the form of wind speed, have received less scrutiny. Recent experimental work has systematically characterized mechanosensory responses of individual neurons within the AL, showing four distinct response patterns, labeled as sustained, transient, biphasic, and offset types; furthermore, these experiments have demonstrated that the distribution of response patterns, as well as the response type of a fixed neuron, can vary with stimulus intensity (wind speed). In this work, we develop a realistic biophysical model of the AL and, using this model, we show that internal AL dynamics are capable of generating the response patterns observed experimentally – namely, we find that response type is determined by the interplay of slow synaptic inhibition, an intrinsic calcium-dependent potassium (SK) current, and stimulus strength, and hence that a heterogeneous distribution of slow inhibition and SK current strength across the AL can lead to the emergence of all four types at a fixed wind speed. Moreover, similar to experiment, we find that the distribution of response patterns across the AL changes with wind speed. Finally, we examine the distribution of response types in the presence of a simulated odor stimulus as well as odor separation by the AL in the presence versus absence of a strong mechanosensory signal.

**Significance Statement:** Odor tracking is a critical for insects, requiring integration of chemosensory (odor) with mechanosensory (wind speed) input. The antennal lobe (AL), the first structure in the insect olfactory pathway, has been extensively studied within the context of odor encoding, but less so in terms of mechanosensory responses. Recent experiments, reported in a companion paper, have systematically studied AL responses to mechanosensory input. In this work, we develop a computational model of the AL to study the network mechanisms that give rise to these empirically observed response patterns. Characterizing the responses of AL neurons to mechanosensory input, and elucidating the network dynamics that give rise to these response patterns, is crucial to understanding sensory integration within the AL.

## 1 Introduction

Insect olfactory navigation – the tracking of an odor source – requires more than simple detection of olfactory stimuli within the environment. Due to air turbuluence, continuous odor filaments emerging from a source rapidly fragment into discontinuous odor strands embedded within clear media, creating a plume structure where short odor strands occur with higher spatial frequency and increasing odor concentration as the source is approached [42, 7, 25, 24]. Thus, odor tracking requires resolving spatiotemporal plume dynamics in a natural environment with turbulent air currents, and integration of chemosensory odor information with mechanosensory wind speed signals is crucial for this task [22]. Accordingly, intermingling of chemosensory and mechanosensory input appears to begin at the periphery; olfactory receptor neurons (ORNs) primarily encode chemosensory signals, but Johnston’s organ and Böhm’s bristles on the antennae are known to guide flight maneuvers through detection of wind speed [29, 16, 35], while other antennal sensilla – including a subtype of trichoid sensilla in the male hawkmoth and sensilla chaetica in the honeybee – actually exhibit bimodality [16, 35, 19, 44].

The antennal lobe (AL) of insects, consisting of excitatory projection neurons (PNs) and inhibitory local neurons (LNs), is the initial processing center within the olfactory pathway for chemosensory information arising from ORNs. While the AL has been extensively studied within the context of odor encoding, mechanosensory encoding within the AL has received less scrutiny, though data suggest that AL neurons (and neurons within the mammalian counterpart, the olfactory bulb) indeed exhibit responses to mechanosensory input in the form of wind buffeting the antennae [13, 6, 15, 2, 40, 39, 28]. The parsing and resolution of the structure of an environmental odor plume may therefore begin at the level of the AL. However, an initial step in understanding the dynamics of sensory integration within the AL is to understand the nature of AL responses to mechanosensory stimulation alone, and the role of AL network dynamics in generating observed response patterns.

Within the honeybee, recordings from AL neurons in response to pure mechanosensory input (i.e., nonscented air puffs) show widespread responses throughout the AL exhibiting a diversity of stimulus-induced firing patterns [39]. As shown in our companion paper [21] (under review), observed response patterns can be categorized as transient (a peak at stimulus inception with rapid decay in spike rate), sustained (elevated firing throughout the duration of the stimulus), biphasic (a peak at stimulus onset with a subsequent rapid decay in firing rate, but with a burst of spikes immediately following termination of the stimulus), and offset (suppressed spiking throughout the duration of the stimulus followed by a spike burst after stimulus termination); additionally, for a fixed AL neuron, response category can change with stimulus intensity (i.e., with increasing wind speed). Elucidating the dynamics underlying the emergence of these various patterned responses is a crucial first step in understanding the integration of mechanosensory and chemosensory input within the AL.

In this study, we modify an AL model developed in prior work [40, 41] in order to assess the emergence of disparate stimulus-induced response patterns through AL network dynamics. We model a mechanosensory stimulus as a broad signal delivered to all AL glomeruli, and we show that a synaptic slow inhibitory current and an intrinsic calcium-dependent potassium (SK) current – when distributed heterogeneously throughout the AL model – can account for the emergence of each of the distinct response categories described above, despite homogeneous stimulus-induced input to all model cells. Furthermore, we demonstrate that the response pattern of a model cell can shift with changing stimulus intensity, and that transitions in response type with increasing wind speed within the model qualitatively match the shifts observed experimentally. We then examine the impact of superimposing an odor stimulus on top of the mechanosensory signal on the nature and distribution of neuronal response types, as well as show that the dynamics of our model AL tend to separate the representations of similar odors.

## 2 Results

The AL model consists of six glomeruli, each containing 10 excitatory projection neurons (PNs) and 6 inhibitory local neurons (LNs), with dense intraglomerular connectivity among PNs and LNs and sparser interglomerular connections mediated solely by LNs. Connections are random but fixed, with connection probabilities dependent on cell type and the intra-vs inter-glomerular nature of the connection. PNs act through fast excitatory (cholinergic) synapses, and are endowed with an intrinsic calcium-dependent potassium (SK) current that is triggered by spiking and serves to curb further firing activity. SK current strength is randomly distributed among PNs. LNs are GABAergic, with synaptic transmission mediated through two types of receptors – fast GABA_*A*_ receptors and slower metabotropic receptors acting over hundreds of ms. In accordance with data showing that mechanosensory responses are widespread within the AL [40], a mechanosensory stimulus (i.e., a nonscented air puff) is modeled as a relatively weak external excitatory current delivered to all glomeruli, while an odor is modeled as a stronger stimulus current restricted to a subset of three glomeruli, with the identity of the odor encoded by the composition of the stimulated glomerular subset (Fig. 1). See *Methods* for details.

**Figure 1.**
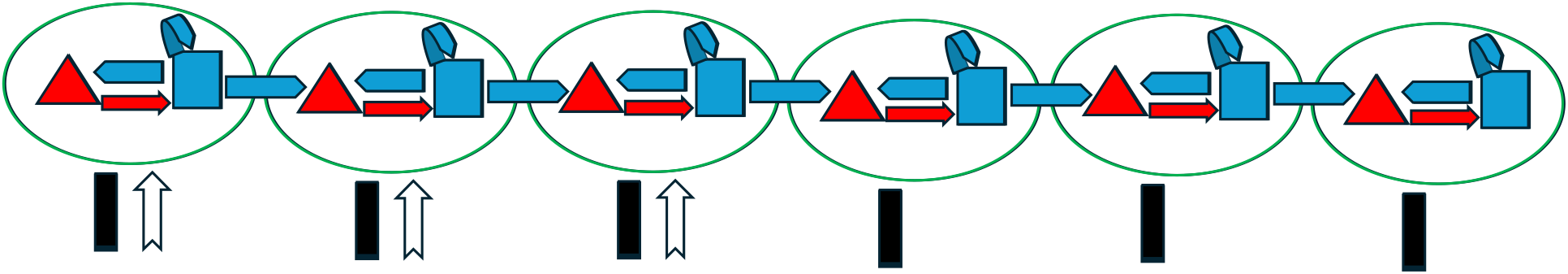
Projection neurons (PNs, red triangle) and local neurons (LNs, blue square) are grouped into six total glomeruli with 10 PNs and 6 LNs per glomerulus (96 total cells). Within a glomerulus, LNs inhibit PNs and other LNs through GABAergic slow and fast inhibition (blue arrow). Further, we model excitatory currents from PNs to LNs (red arrow), however, due to a lack of experimental evidence, our network contains no PN-to-PN synapses. Interglomerular connectivity is mediated by LN to PN synapses linking all glomeruli. Mechanosensory input (black bar) is modeled as an excitatory stimulus to all PNs and LNs in all glomeruli. Odor stimulus (white arrow) is represented as a strong excitatory current to all cells within a subset of three glomeruli.

### 2.1 Network Dynamics and Response Patterns

Experiments described in our companion paper [21] (under review) show that during mechanosensory stimulation, AL cells can exhibit several distinct slow response patterns, which can be grouped into sustained, transient, biphasic, and offset types. Our model successfully replicates each of these responses, as illustrated by the histograms in Fig. 2. Sustained responses are characterized by an elevated firing rate during the entirety of the stimulus (Fig. 2, top row), while transient responses have an initial high intensity burst at the inception of the stimulus (for the first ∼200 ms) followed by a rapid decrease in firing rate back to or slightly above background for the remainder of the stimulus (Fig. 2, second row). Biphasic PNs tend to spike initially at stimulus onset, then display a markedly lower firing rate for the remainder of the stimulus, with a renewed burst of spiking activity after stimulus termination (Fig. 2, third row). Offset PNs display suppressed activity during the stimulus, but exhibit a burst of activity following stimulus termination (Fig. 2, bottom row). See *Methods* for details of the criteria employed to classify response type within the model.

**Figure 2.**
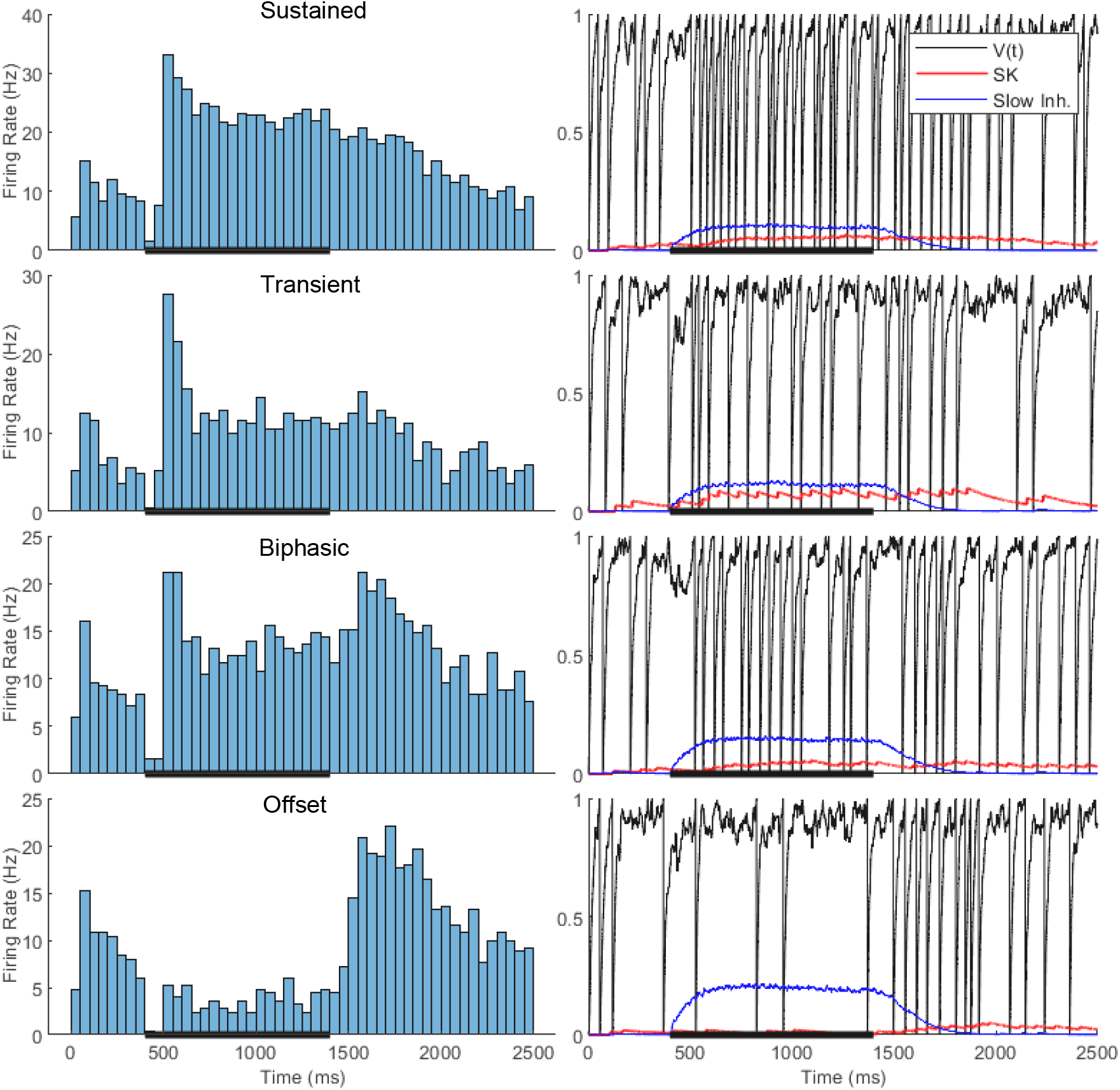
Firing rates averaged over 50 trials (left) and the plotted membrane potential, slow inhibition, and SK current (right) of sample network PNs in response to mechanosensory stimulus pulse alone. Each PN was chosen randomly from the group of PNs classified in a given response type. From top to bottom, response types are classified as sustained, transient, biphasic, and offset. Black bar along the horizontal axis represents one-second stimulus duration; different trials correspond to different representations of the Poisson noise received by each cell, with network connectivity and parameters fixed across trials.

The disparate response patterns between network PNs prompts the question: what components of the model give rise to these emergent behaviors? We find that variations in two key network elements contribute to the generation of distinct PN response patterns: (1) the level of inhibition (notably slow inhibition) received by a PN; (2) the strength of the SK current within a PN. This is supported by Fig. 3 – the left panel of Fig. 3 shows that PN response types display a strong tendency to cluster within the two-dimensional parameter space defined by the number of presynaptic LNs (a proxy for the level of fast and slow inhibition received) and SK current strength. Fig. 3 (middle) shows that without fast GABA_*A*_ receptors, the temporal structure of response patterns remains largely unchanged across network PNs (though we find that firing rates –especially during stimulus onset– tend to increase somewhat, which can alter the response type of some individual PNs); in contrast, without slow inhibition (Fig. 3, right), all PNs in the network shift to either a sustained or transient pattern. This suggests that, within the model, while fast inhibition may augment the suppressive effect of the slow inhibitory current, slow inhibition plays the more prominent role in generating the diversity of response types. We therefore focus on slow inhibition and the SK current as the primary determinants of PN response patterns.

**Figure 3.**
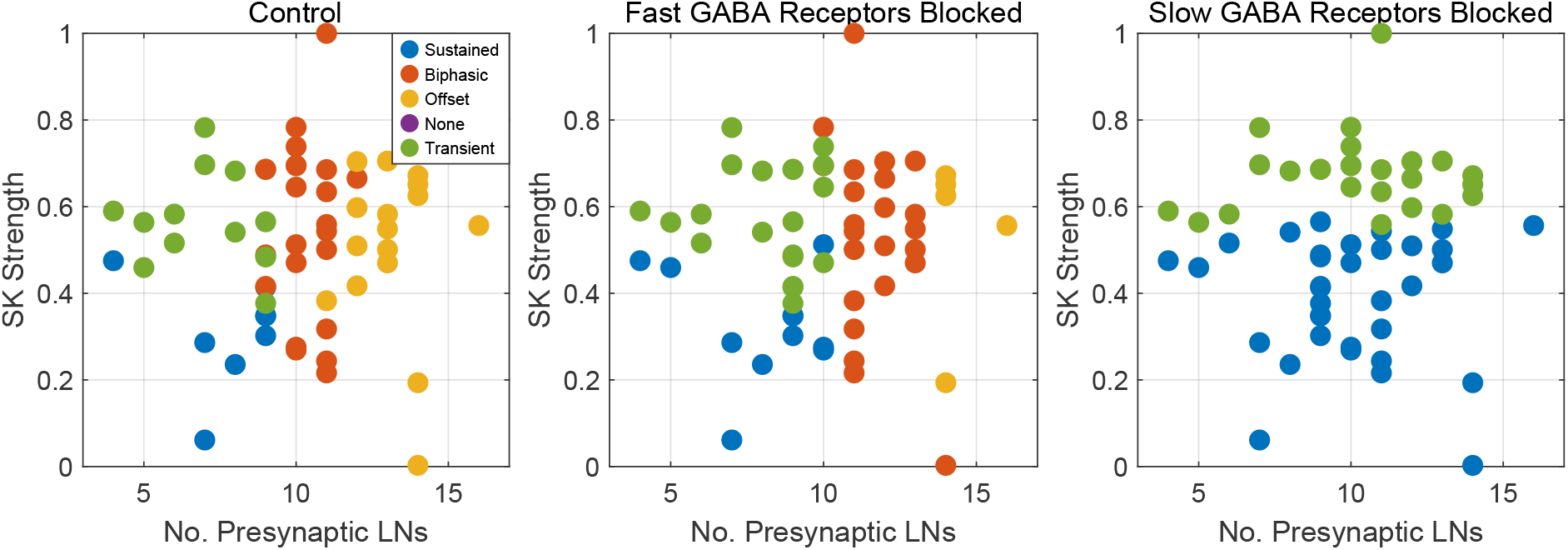
SK current strength vs. presynaptic LN connections in AL network PNs grouped by response type with fast and slow inhibition (left), without fast inhibition (middle), and without slow inhibition (right). Note that removing slow inhibition makes all PNs sustained or transient responders. Response types were determined from firing rates averaged over 50 trials.

In Fig. 3 (left), we see that network PNs with a higher than average number of presynaptic LN connections (greater than 11) tend to have an offset response to the stimulus, an intermediate number of LN connections (9-11) is correlated with a biphasic response pattern, and PNs with lower than average LN connections (less than 9) tend to be transient or sustained. Moreover, we see that, for PNs with a lower number of presynaptic LNs, transient versus sustained patterns cluster by SK strength – transient PNs have a stronger SK current than sustained PNs. Fig. 4, in which we construct four specialized networks to generate a single response pattern in all network PNs, further supports this clustering structure. Severing global LN→PN synapses and decreasing the SK current strength across the network yields sustained PN responses (Fig. 4A), while increasing the potency of the SK current (while leaving LN→PN synapses blocked) produces transient response types (Fig. 4B). Additionally, if SK current strengths remain at standard values, manually setting the number of LN→PN connections slightly above the average for the standard network causes PNs to become predominantly biphasic (Fig. 4C), while setting this number at a high value yields the offset response pattern (Fig. 4D).

**Figure 4.**
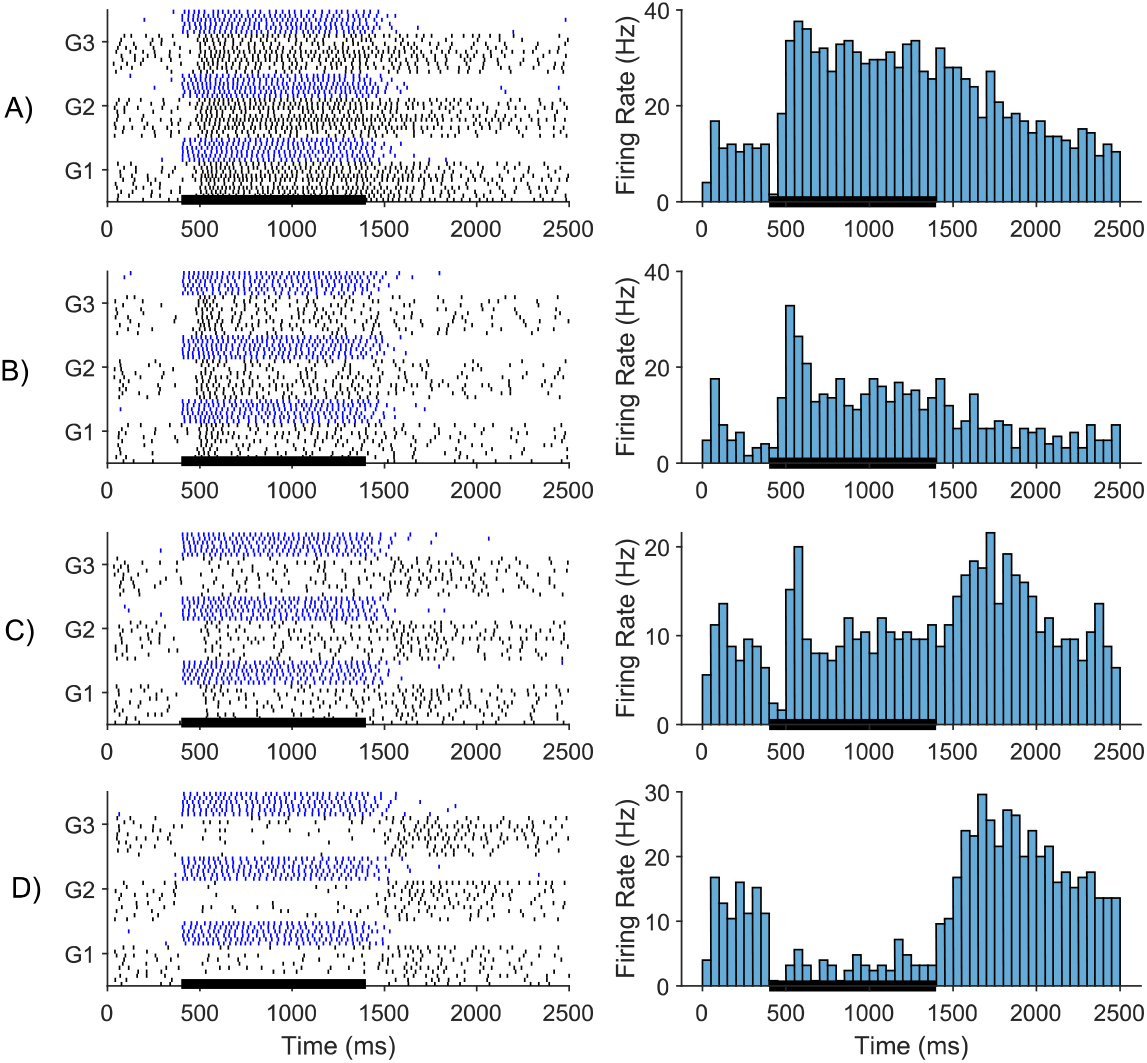
Raster plots of the first three glomeruli and a representative PN from four specialized networks for generating each response type. Sample PNs were chosen randomly from the network. PNs are in black, LNs are in blue. (A) Sustained: blocked interglomerular LN-to-PN synapses. Decreased the mean SK strength of network PNs. (B) Transient: blocked interglomerular LN-to-PN synapses. Increased the mean SK strength of network PNs. (C) Biphasic: manually set the number of presynaptic LN connections to 11 for each PN. Did not change SK strength. (D) Offset: increased the mean number of LN connections to 14. Did not change SK strength.

To ensure that the results in Fig. 4 can be properly interpreted within the context of individual PN dynamics, rather than as being an emergent consequence of the global parameter changes employed in the simulations, we take a PN from the standard network and systematically vary the strength of its SK current and slow inhibitory synapses (Fig. 5). Across each row, Fig. 5 shows that, consistent with the results presented in Figs. 3 and 4, increasing slow inhibitory strength from low to high transforms the response of this PN from sustained/transient to biphasic to offset. At lower values of slow inhibition (Fig. 5, columns one and two), we see that SK strength modulates the difference between transient and sustained response patterns (the stronger the SK current, the more transient the response), also consistent with Figs. 3 and 4. However, we note that the distinction in SK strength between transient and sustained response types is dependent on slow inhibition as well: the response pattern is more transient at weaker SK strengths in column two versus column one of Fig. 5.

**Figure 5.**
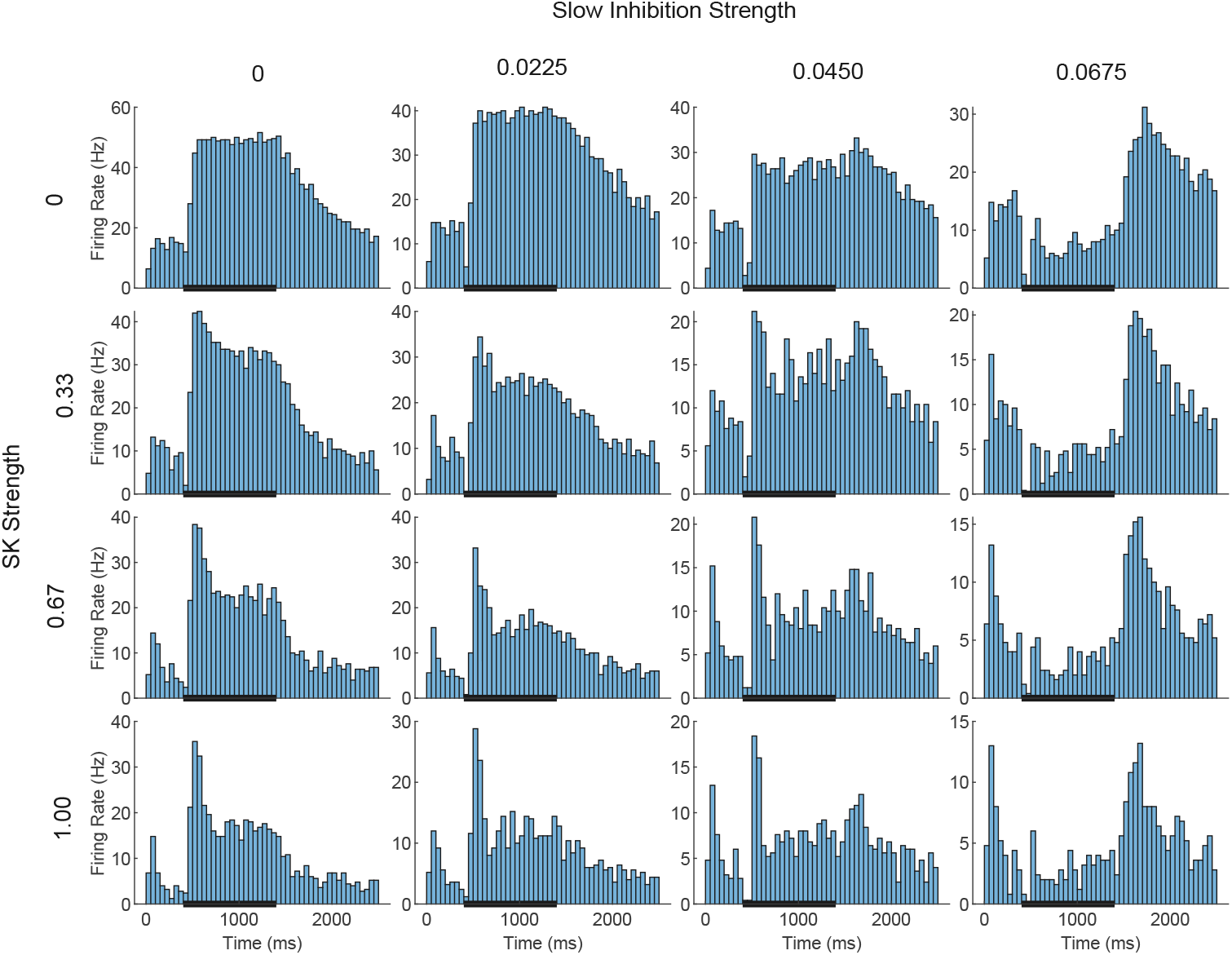
We selected a network PN with 10 presynaptic LNs (the network average LN-to-PN connections) and systematically varied the strength of its SK current and slow inhibitory synapses. Across the rows, slow inhibition increases in increments of 0.0225. Down the columns, SK strength increases in increments of 0.33. The standard strength of an LN-to-PN slow inhibitory synapse is 0.0475, the average number of presynaptic LNs for a PN is 10, and for a standard PN SK current strength is drawn from a Gaussian distribution with mean equal to 0.5 and a standard deviation of 0.2. Firing rates were calculated as the average over 50 trials.

A question then naturally arises: what are the dynamics underlying the modulation of response type by slow inhibition and the SK current? The sustained response pattern is the simplest. In this case, due to the low number of presynaptic LNs, inhibition is not strong enough to prevent PN spiking at stimulus onset, and the weak SK current allows for continuous firing during the stimulus window. Consequently, the stimulus-induced firing rate of sustained responders is the greatest out of all four response patterns. When a PN’s SK current strength is high and synaptic inhibition remains low, the stimulus evokes an initial high-intensity burst, but the resulting SK activation dampens continued firing, producing a transient response. Furthermore, the strongly active SK current prevents a notable spike burst following stimulus offset (as evidenced by the fact that transient response patterns can be obtained with slow inhibition blocked – see Fig. 5, bottom left panel). As the slow inhibitory current to a PN becomes stronger, PN stimulus-induced firing rates decrease, and SK strength is less relevant in shaping response patterns. Biphasic responders receive intermediate levels of slow inhibition. This tempers initial PN firing, providing less of an initial trigger for the SK current, and full activation of slow inhibition over a few hundred ms suppresses firing throughout the remainder of the stimulus, which further reduces SK current activation. Following stimulus offset, the time scale of decay of mechanosensory input to PNs is longer than that of the slow inhibitory current received by PNs; coupled with the lack of significant SK current activation during the stimulus, this allows PNs to fire briefly after the stimulus pulse concludes. Offset PNs have the highest number of presynaptic LNs and receive the most slow (and fast) inhibition. Thus, firing is suppressed in these PNs throughout the entire duration of the stimulus, which again leads to a lack of SK current activation; following stimulus offset, the temporal asymmetry between the decay time of stimulus current versus slow inhibition to PNs, along with a lack of SK current, triggers a post-stimulus spike burst. Since the stimulus current decays following the conclusion of the stimulus, offset responders have the lowest firing rates compared to the other response types.

These results represents two experimentally testable predictions of the model – firstly, the model predicts that, within the AL, LN-to-PN connectivity is heterogeneous across the network and the level of stimulus-induced slow inhibition received by a PN determines its response type to mechanosensory input. Second, the model predicts that the SK current varies across PNs and the strength of a PN’s SK current differentiates between transient and sustained responses to mechanosensory stimulation.

### 2.2 Effect of Increasing Stimulus Intensity on Response Patterns

Experimental evidence shown in our companion paper [21] (under review) has demonstrated that when wind speed is increased, the response type of an individual PNs can shift; notably, the sustained or transient responses tend to dominate at higher wind speeds. We therefore examine model behavior as mechanosensory stimulus strength is varied (Fig. 6). In Tables 1 and 2, we see that the experimentally validated changes in response pattern with wind speed are captured within the model. As wind speed in increased, PNs with an initial offset response at a lower wind speed tend to transform into biphasic responders at higher wind speeds, while biphasic responders tend to become transient responders, transient responders eventually become sustained, and sustained responders tend to remain unchanged.

**Table 1.** Proportion of each response pattern transforming between 2.0 and 4.0 spikes/sec mechanosensory input rate.

|  |  | 4.0 Hz |  |  |  |
| --- | --- | --- | --- | --- | --- |
|  |  | Sustained | Transient | Biphasic | Offset |
| 2.0 Hz | Sustained | 62.50% | 37.50% | 0.00% | 0.00% |
|  | Transient | 0.00% | 100.00% | 0.00% | 0.00% |
|  | Biphasic | 4.34% | 17.40% | 78.26% | 0.00% |
|  | Offset | 0.00% | 0.00% | 70.59% | 29.41% |

**Table 2.** Proportion of each response pattern transforming between 4.0 and 6.0 spikes/sec mechanosensory input rate.

|  |  | 6.0 Hz |  |  |  |
| --- | --- | --- | --- | --- | --- |
|  |  | Sustained | Transient | Biphasic | Offset |
| 4.0 Hz | Sustained | 100.00% | 0.00% | 0.00% | 0.00% |
|  | Transient | 21.05% | 78.95% | 0.00% | 0.00% |
|  | Biphasic | 3.33% | 30.00% | 66.67% | 0.00% |
|  | Offset | 0.00% | 0.00% | 60.00% | 40.00% |

**Figure 6.**
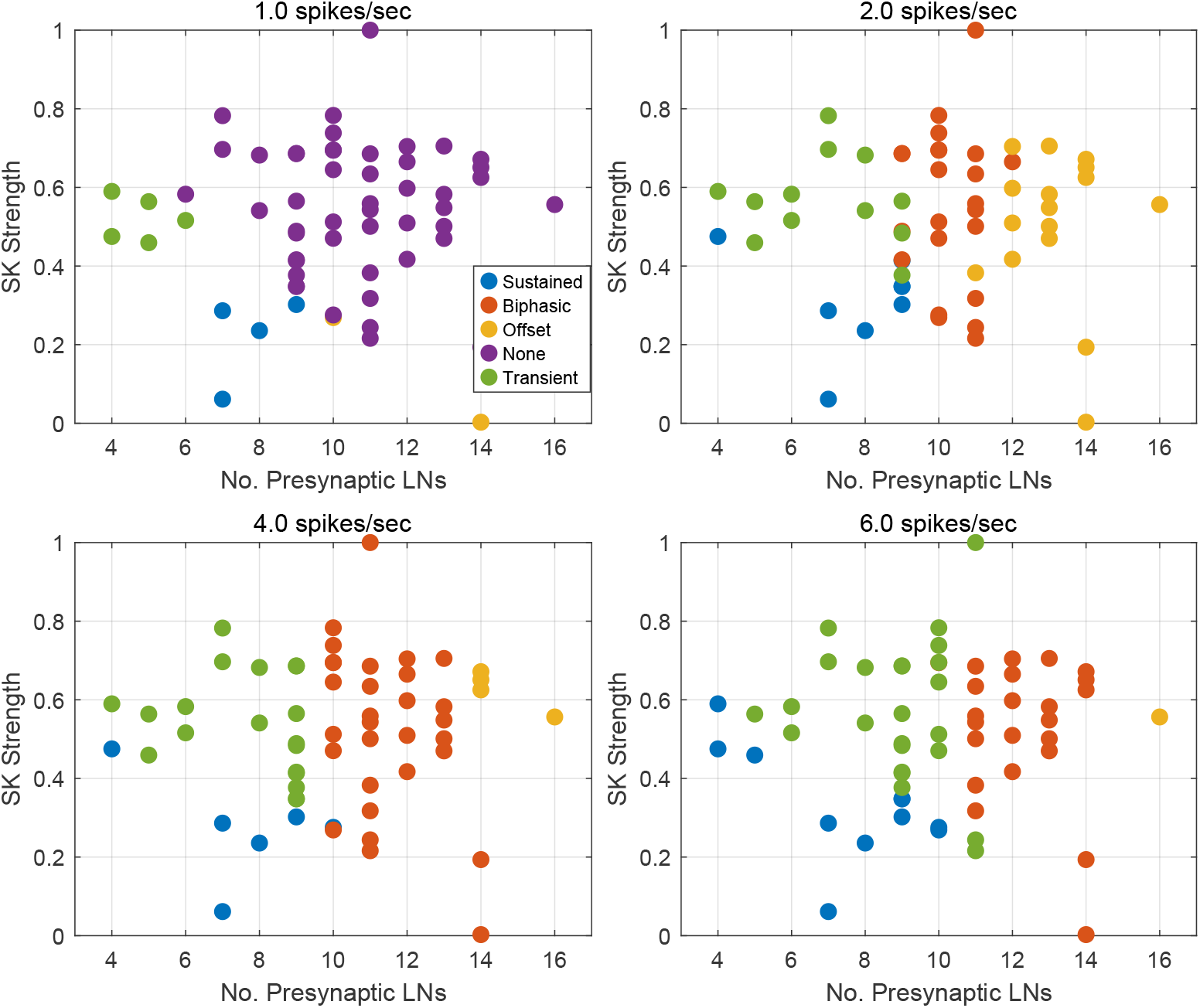
Proportion of PNs switching response between 2.0 spikes/sec and 4.0 spikes/sec and 4.0 spikes/sec and 6.0 spikes/sec, grouped by response at lower wind input (top). Plotted SK current strength vs. presynaptic LN connections in network PNs grouped by response type across increasing mechanosensory input rate (bottom) As mechanosensory input increases, offset responses become biphasic, biphasic become mainly transient, and transient eventually become sustained. Responses were determined from firing rates averaged over 50 trials.

The network mechanisms behind the variation in PN response patterns across mechanosensory input strengths can be gleaned from Fig. 6. At the lowest stimulus strength of 1.0 spikes/sec, only PNs with weak SK currents or a low number of presynaptic LN connections – i.e., PNs receiving little intrinsic or synaptic inhibition – exhibit noticeable responses above background, and, consistent with the results of the previous section, the few PNs that respond show transient or sustained spiking patterns. At all other stimulus intensities, every PN responds, and we can see the familiar clustering of response patterns in the two dimensional parameter space of SK current strength and number of presynaptic LNs.

From 2.0 spikes/sec to higher stimulus strengths, Fig. 6 shows that the average number of presynaptic LNs for PNs of each response type increases. For instance, offset responders at 6.0 spikes/sec have a higher number of presynaptic LNs than offset responders at 2.0 spikes/sec. This suggests that as the mechanosensory input to PNs increases, the interplay between stimulus strength and slow inhibition causes the variation in response patterns among network PNs. If a PN is an offset responder at low wind speed, then at a higher wind speed the increased stimulus-induced input can overwhelm the incoming pooled inhibition at the inception of the stimulus, inducing onset firing. Since an offset responder has a relatively high number of presynaptic LNs, the resulting onset spiking is quickly suppressed as slow inhibition fully activates over a few hundred ms, which in turn reduces SK current activation, rendering the SK current unable to prevent post-stimulus spiking. Hence, for moderate increases in mechanosensory input rates (e.g. 2.0 to 4.0 or 4.0 to 6.0 spikes/sec), offset responses tend to transform into biphasic responses.

The transformation of biphasic responses into transient responses is similar. Increased stimulus-induced input induces greater firing rates at stimulus onset, activating the SK current; however, the increased level of stimulus current ensures that, after an onset spike burst, spiking is reduced due to slow inhibition (since these cells have a considerable number of presynaptic LNs) but still sufficient to retain substantial SK current activation, preventing rebound firing post-stimulus. Transient responses that vary as wind speed increases only become sustained. We observe this transformation primarily in PNs that are transient at low wind speed and hence have few presynaptic LNs but high SK current strengths. As mechanosensory input increases, the stimulus strength overwhelms the high SK current strength, and in conjunction with relatively weak incoming synaptic inhibition, drives continuous firing throughout the duration of the stimulus. We do see a proportion of sustained responders become transient from 2.0 spikes/sec to 4.0 spikes/sec. This may be due to slow inhibition from increased LN activity at a higher input rate pushing a PN that is ‘in between’ a transient and sustained responder into a transient response mode. Overall, though, sustained responses tend to stay sustained across mechanosensory input rates.

### 2.3 Odor-Induced Dynamics

Many important odorants in a honey bee’s natural environment are composed of mixtures of structurally similar molecules that bind to overlapping groups of ORNs [17, 30]. Also, a given odorant is unlikely to be encountered in isolation – a bee receives olfactory information from many sources simultaneously. Thus, it is an essential processing function of the AL to disentangle similar chemical signals by converting similar ORN input patterns into distinct neural representations. Mechanosensory input within the AL, which is not odor specific and has been experimentally shown to be a relatively global stimulus [40], may further confound this effort. In this section, we test the model’s response to a single odor – simulated as a pulse of excitatory input to all cells in a subset of glomeruli (glomeruli 1-3) – in addition to mechanosensory stimulation. Then, we simulate two structurally similar odors, with and without mechanosensory input (i.e., at high and low speed), and quantify the ability of the AL network to separate their representations.

Because the odor is a focal stimulus only to glomeruli 1-3, the LN mediated inhibition a PN receives is no longer directly proportional to the number of presynaptic LN connections, since LNs within glomeruli 1-3 are more strongly activated than those within glomeruli 4-6. Additionally, PNs within glomeruli 1-3 receive considerably more external excitation than those in glomeruli 4-6. Thus, the odor differentially affects response patterns in glomeruli 1-3 versus 4-6, as shown in Fig. 7. The PNs in glomeruli 1-3 have mainly transient response patterns across the first three mechanosensory input rates (Fig. 7, top four panels). The strength of the additive wind and odor stimulus delivered to PNs in these glomeruli ensures onset spiking is always present (precluding the existence of the offset response pattern), but following an initial spike burst the local LN→PN connections aid the SK currents in suppressing continued firing. Those PNs with fewer presynaptic LNs and weaker SK currents tend to be sustained, and those with more presynaptic LNs tend to be biphasic because the initial firing burst is subsequently tempered by slow inhibition, thus yielding insufficient activation of the SK current that is necessary to prevent offset spiking. At the highest mechanosensory stimulation strength, PNs in glomeruli 1-3 receive enough excitation to ensure onset spiking, but the highly activated local LN→PN connections quickly suppress PN firing to below background rate. This results in inactive SK currents across PNs, allowing for firing post-stimulus.

**Figure 7.**
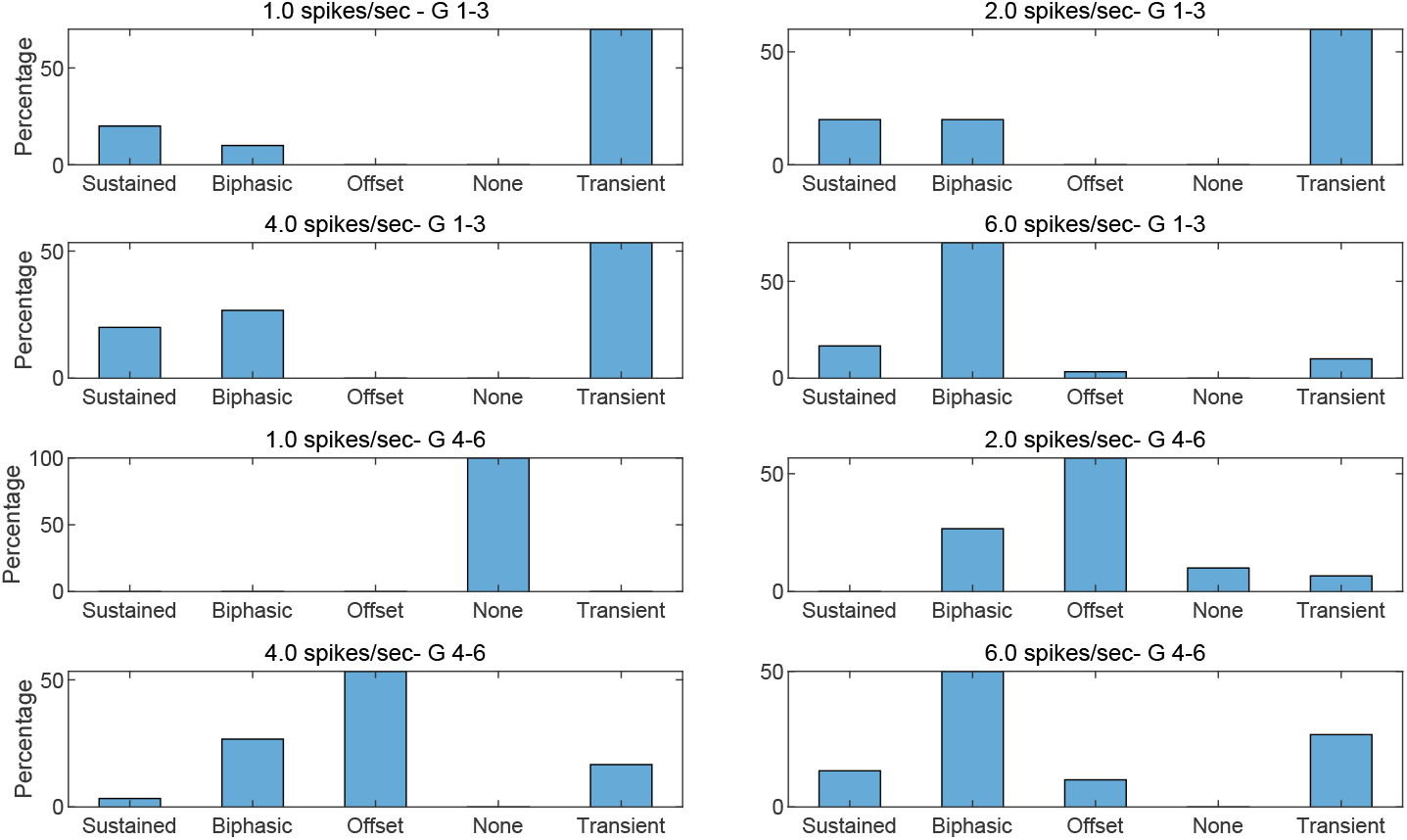
Distribution of PN responses with odor stimulus across increasing mechanosensory input. Odor is simulated as a one second pulse to only glomeruli 1-3 (top four panels). Glomeruli 4-6 (bottom four panels) receive mechanosensory input alone. The increased inhibition from glomeruli 1-3 affects the responses of PNs in glomeruli 4-6. Responses were determined from firing rates averaged over 50 trials.

The responses of PNs in glomeruli 4-6 (Fig. 7, bottom four panels), which receive no odor-induced excitation, follow the dynamics of response pattern shifts with changing wind speed discussed in the previous section (Fig. 6). However, because of the strengthened global inhibition from LNs in glomeruli 1-3, we see a higher proportion of offset responders among these PNs. Additionally, instead of only being dependent on the total number of presynaptic LNs, the glomerular location of LN connections is also a factor in determining a PN’s response type. For instance, when comparing two PNs from glomeruli 4-6 with the same number of presynaptic LNs, one PN could have many LN connections from glomeruli 1-3 and be offset, while the other PN has few presynaptic LNs from glomeruli 1-3 and be sustained. The difference in the response dynamics between these two PNs is due the highly activated LNs in glomeruli 1-3.

We further assess the ability of the network to separate the representations of similar odors. Two structurally similar odors are simulated separately: odor 1 to glomeruli 1-3, odor 2 to glomeruli 2-4. For each odor, we measure the Euclidean distance between the input vector (the vector of incoming stimulus current to all PNs within the network) and output vector (the trial-averaged vector of PN firing rates in response to odor presentation) in 50 ms increments. We note that, in order to avoid effects of vector magnitude on Euclidean distance, we normalize input and output vectors so that vector components range between 0 and 1 (see *Methods* for details).

Fig. 8 shows the separation between the input and output vectors for each odor at low (Fig. 8, left) and high (Fig. 8, right) wind speed. In both cases, the model successfully separates odor representations above the baseline separation of input vectors. However, the odor-only stimulus scenario exhibits the greatest separation of odor representations. This matches intuitive expectations – wind stimulus is nonspecific and independent of odor identity, and thus increases the uniformity of the input current across PNs, confounding odor identity information.

**Figure 8.**
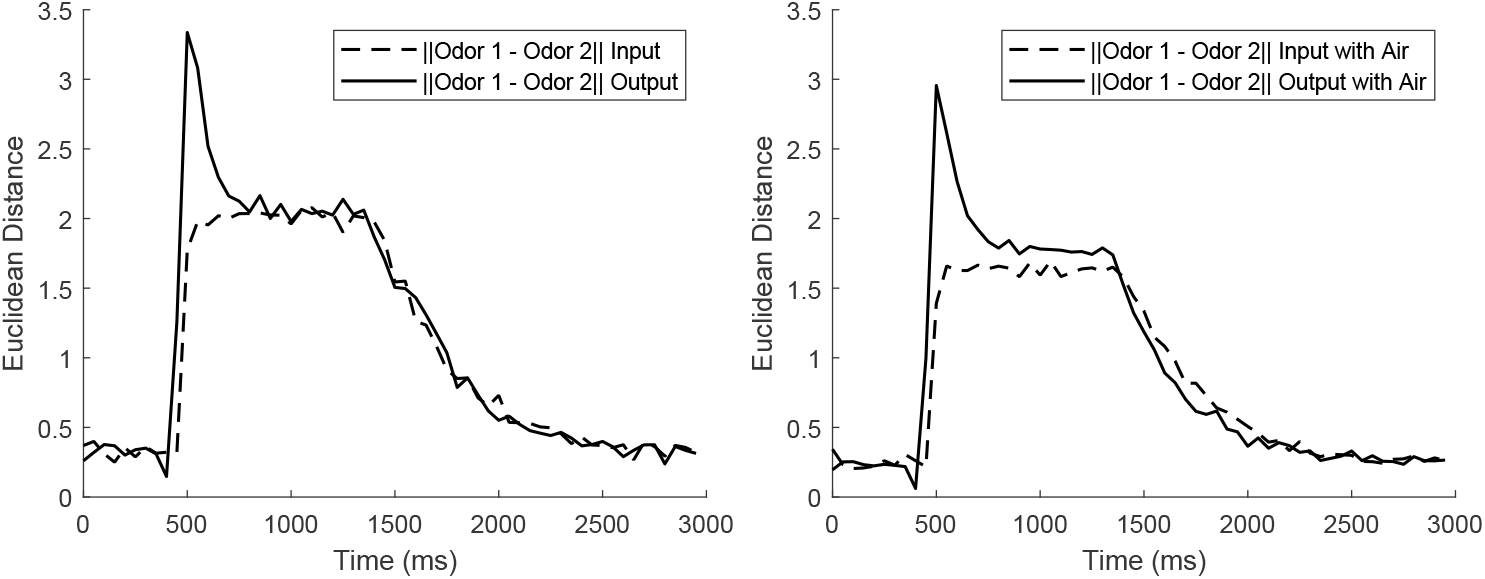
We simulated two similar odors (odor 1: glomeruli 1-3; odor 2: glomeruli 2-4) separately and measured the Euclidean distance between the 60-dimensional PN input and output vectors in 50 ms increments (see Methods for details). On the left, distances were measured without mechanosensory input; on the right, mechanosensory stimulus was included.

We also see that, both with (Fig. 8, right) and without (Fig. 8, left) mechanosensory input, odor separation by the model is greatest in the first few hundred milliseconds of stimulus presentation. Within this initial interval, little slow inhibition has accumulated for most PNs, leading to high-magnitude spike bursts, while for other PNs (namely, offset responders), LN inhibition suppresses firing rates below background; notably, even within a single glomerulus with a common input pattern across both odors, the identity of PNs that exhibit a high-amplitude spiking response versus those that are suppressed may differ between odor 1 and odor 2. Hence, the most pronounced differences in PN firing rates between odor 1 and odor 2 are seen in this period, resulting in increased odor separation. Beyond this initial period, as the nascent slow inhibitory current grows, PN firing decreases across the AL and odor separation by the model diminishes and approaches the separation of input vectors. This is in concordance with empirical evidence showing that odor identification tends to occur within the first few hundred ms of an odor encounter [18].

## 3 Discussion

In this work, using a biophysically detailed computational model, we investigate the network components driving the distinct mechanosensory-evoked response patterns across PNs in the honey bee AL. Our results suggest that a combination of LN→PN slow inhibition and the differences in SK current strengths across PNs causes the disparate PN response dynamics within the model. We then use our model to dissect the internal mechanisms behind the effects of increasing mechanosensory input on PN dynamics. Our modeling matches recent experimental data reported in our companion paper [21] (under review), and we show that the interaction between mechanosensory stimulus strength and LN→PN slow inhibition is responsible for the variation of PN responses across increasing wind speeds. Thus, we show that varying response patterns across PNs for a fixed stimulus, and in a fixed PN across varying stimulus intensities, can be explained by internal network dynamics rather than requiring patterned ORN input.

Finally, we simulate both olfactory and mechanosensory input within our model. The focal nature of olfactory input alters the uniformity of the LN→PN slow inhibitory current across glomeruli, causing varying PN responses dependent on the glomerular location of a PN and its presynaptic LNs. Lastly, our simulations suggest that, while the representations of similar odors diverge due to AL dynamics, mechanosensory input decreases the distance between structurally similar odors within the AL coding space.

### 3.1 Dynamics of Patterned Responses

While responses to mechanosensory input have been observed repeatedly within the AL in empirical studies [13, 6, 15, 2, 40, 39, 28], the precise source of the mechanosensory input to the AL that elicits such responses has not been pinpointed experimentally. Empirical and theoretical studies are suggestive of several possibilities for the transmission of mechanosensory input to the AL, including the existence of ORN responses to mechanosensory stimulation [11, 8], or ephaptic coupling within the olfactory nerve causing olfactory fibers to be stimulated by the mechanosensory fibers [4], or direct or indirect transmission to the AL by central structures – namely, the antennal mechanosensory and motor center (AMMC) – that process mechanosensory input [23, 1].

Regardless of the source of mechanosensory input to the AL, the existence of diverse response patterns, and the shifts in pattern type with changes in stimulus intensity, may arise through a mechanism external to the AL or through internal AL dynamics (or a combination of both). It is possible that mechanosensory input is spatially nonuniform throughout the AL and exhibits diverse, heterogeneously distributed response patterns that shift with stimulus intensity, and that AL neurons simply inherit their response characteristics from the input. In constrast, mechanosensory input to the AL may be more uniform and lack patterned response structures, and hence the diversity of response patterns (along with the changes in response patterns with stimulus strength) arise from internal AL dynamics. In this study, we show that patterned, heterogeneously distributed input is not necessary to generate the diverse response patterns that shift with stimulus intensity observed experimentally [21] (under review), and that these features can arise through relatively simple and realistic AL network components – namely, slow synaptic inhibition coupled with an intrinsic SK current.

### 3.2 Biological Implications

The results presented in this work raise the question: what is the biological significance of the diversity of neuronal responses to mechanosensory input in the honey bee AL? The approximately 800 PNs in the honey bee AL innervate the mushroom body and synapse onto tens of thousands of Kenyon cells [10, 37]. The network of Kenyon cells in the mushroom body is crucial for stimulus recognition and olfactory memory storage [27]. Thus, the significance of PN response pattern variation, as suggested by our results in this paper, may be that it separates the representations of similar odors within the AL, making them easier to distinguish by downstream Kenyon cells. This leads to less overlap in the representation of molecularly similar odors at the Kenyon cell level (compared to at the ORN level), entailing less cross-odor interference during learning and memory processes within the mushroom body.

Honey bees are social, foraging insects that must explore the landscape for colony success. Bees in the hive receive information about pollen and nectar sources from ‘scout’ bees via the waggle dance [43]. When bees lack access to waggle dance information from nest mates, they must use their navigational prowess to forage, possibly miles away from the hive [33]. To do this, bees use an optimal flight pattern known as Lévy flight where they surge in a straight path upwind in the direction of an odor source, while incorporating random directional changes to increase the chance of encountering the next odor target [31, 32]. Detecting wind speed allows bees to tune their Lévy flight path to real-time conditions by modulating flight muscles and the metabolic cost of flying. The PN response patterns to mechanosensory input presented in this paper, specifically the predictable variation in PN responses to increasing wind speed, may allow the bee to integrate wind speed information within the AL to assist in odor tracking while foraging.

Another aspect of insect behavior that may be affected by wind speed is active sampling. Air turbulence has been shown to fragment an odor plume into concentrated packets separated by pockets of low odor concentration [24, 26]. Insects sample the odor plume by flicking their antenna towards locations of higher odor concentration [14]. Thus, the neural processing of wind velocity within the AL may help insects optimize flicking patterns to better encode the spatial and temporal structure of odor plumes.

While odor tracking is important for survival, honey bees also process many odors in low wind environments, such as in the hive. Honey bees use pheromone-based communication in many different aspects of colony ecology, from regulating reproduction to promoting social cohesion [5, 36]. Hence, increased separability of odors in low wind environments, as suggested by our model, may aid in the processing of pheromone mixtures by the olfactory system within the hive. This is supportive of the odor discrimination versus source tracking hypothesis proposed in prior work [40, 41, 28]: mechanosensory input enhances the ability of AL activity to track and home in on an odor source while impairing fine odor discrimination, consistent with the former task usually being more important in a high mechanosensory input context (e.g., when the insect is flying through the air to forage) and the latter task having higher relative importance in lower mechanosensory input environments (e.g., when the insect is sitting relatively still on a flower). Overall, the integration of mechanosensory and olfactory input within the AL likely has a multifaceted effect upon insect behavior. Further experimental and theoretical work will be necessary for a more thorough understanding of the dynamics underlying this form of sensory integration.

## 4 Methods

We developed a firing model of the honey bee antennal lobe (AL) that obtains realistically complex behavior but maintains a level of simplicity that is suitable for investigation of basic network components. Our model was primarily based on previous computational work on the moth AL [20, 40, 41]. In the following sections, we elaborate on the mathematical basis and structure of our network, as well as methods used to analyze network dynamics.

### 4.1 Individual neuron model

The honey bee AL model includes two categories of neurons: excitatory cholinergic projection neurons (PN) and inhibitory GABAergic local neurons (LN) [3, 9, 34]. The membrane potentials of the *j*^*th*^ PN, denoted 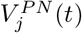 (where *t* is the simulation time measured in milliseconds), and of the *j*^*th*^ LN, denoted 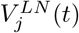, were modeled using the following set of integrate-and-fire ordinary differential equations:

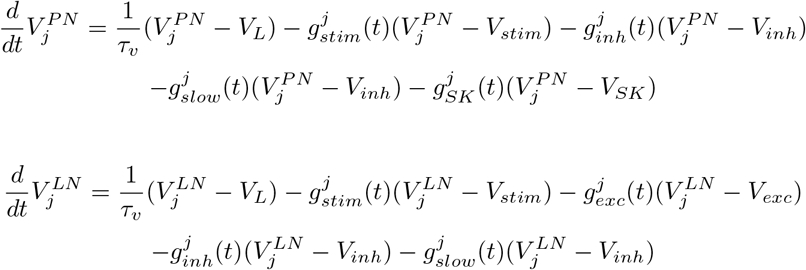

PNs receive stimulus-based excitatory input and slow and fast inhibition from other LNs in the network. Although SK currents have not been identified experimentally, PNs exhibit calcium-activated potassium currents, which may include SK-type channels [12]. LNs receive stimulus-based excitatory input, excitatory input from other PNs, and slow and fast inhibition from other LNs. The parameters: 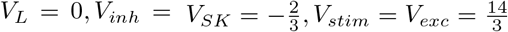 denote the nondimensional reverse potentials for leak, inhibition (including SK mediated), and excitation currents, respectively. The leakage timescale, *τ*_*V*_, was set to 20 ms. When a neuron’s membrane potential reaches the threshold value *V*_*thresh*_ = 1, a spike was recorded and the membrane potential was reset to zero for a refractory period of *τ*_*ref*_ = 2 ms. The reduced integrate-and-fire differential equations were based on a model previously developed in the literature [38].

The term 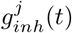 denotes the synaptic conductance of neuron *j* to fast GABA_*A*_ inhibitory input from LNs. This was modeled as follows:

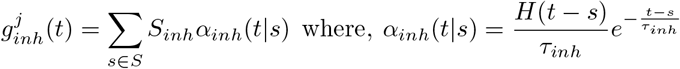

where *H*(*t*) is the standard heavy-side step function:

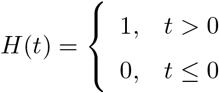

In this model, *S* represents the set of all spike times of all presynaptic LNs to neuron *j*. The term *S*_*inh*_ is the coupling strength of an LN to neuron *j*. If neuron *j* is a PN, *S*_*inh*_ = 0.007; if neuron *j* is an LN *S*_*inh*_ = 0.015. Lastly, *α*_*inh*_(*t*|*s*) is a function with an instantaneous rise time and an exponential decay time set at *τ*_*inh*_ = 2 ms for both LNs and PNs.

The conductance terms for excitation, 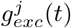, slow inhibition, 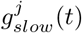, and stimulus input, 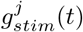, were modeled in the same way:

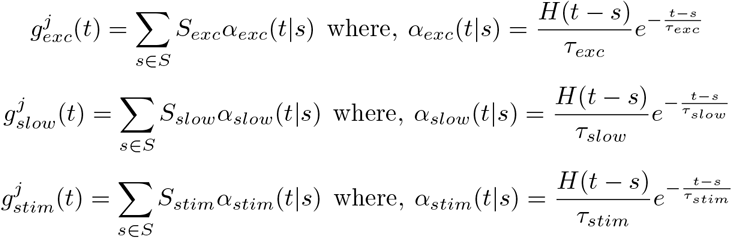

For 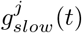, *S* again denotes the set of all spike times of presynaptic LNs to neuron *j*. For 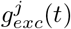, *S* is the set of all spike times from PNs synapsed to neuron *j*. For 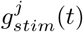, *S* is the set of all external spikes to neuron *j*, that is, background noise, odor, and mechanosensory inputs represented as a Poisson processes (see *Stimulus Modeling* for more details). If neuron *j* was a PN, *S*_*slow*_ = 0.0475 and *S*_*stim*_ = 0.0040 (note: PNs are modeled with no incoming PN-to-PN connections). If neuron *j* was an LN, *S*_*exc*_ = 0.006 *S*_*slow*_ = 0.04, and *S*_*stim*_ = 0.0032. The values for the decay timescales are as follows: *τ*_*exc*_ = *τ*_*stim*_ = 2 ms, *τ*_*slow*_ = 100 ms for PNs, and *τ*_*slow*_ = 450 ms for LNs.

The SK current is an intrinsic inhibitory current unique to PNs in the model. Upon spiking, the SK current will activate to inhibit continued spiking. Instead of instantaneous activation when a PN fires, the SK current was modeled with a sigmoidal rise time and exponential decay. The longer rise time allows PNs to potentially fire multiple times before being suppressed by the activation of the SK current. The conductance of the SK current for PN *j* was modeled as follows:

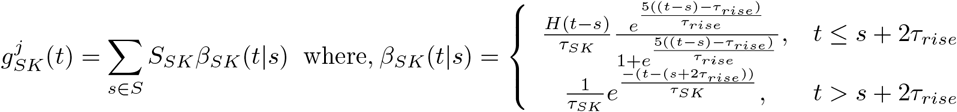

S represents the set of all spike times of PN *j*. The rise time was set to *τ*_*rise*_ = 25 ms and the decay time was *τ*_*SK*_ = 250 ms. *S*_*SK*_, the strength of the SK current, was randomly determined from a normal distribution with *µ* = 0.5 and *σ* = 0.2. Any negative value of *S*_*SK*_ was manually set to zero.

### 4.2 Network design

The AL network was organized into 6 glomeruli; each glomeruli contained 10 PNs and 6 LNs. The connectivity between cells was fixed, randomly determined, and neuron type dependent. The relatively dense connections within a glomeruli were defined by the following probabilities : LN→ PN = 0.65, LN→ LN = 0.75, and PN→ LN = 0.5. No direct PN→PN connections were included in the model due to lack of experimental evidence. Connectivity was sparser between glomeruli: we chose to model only LN→PN synapses with a probability of 0.23. Fig. 1 depicts a simple schematic of the network architecture.

Note that the exact parameters provided are not necessary for producing realistic network behavior. For example, in our model, we primarily used slow inhibition to suppress PN firing during stimulus presentation. However, weakening slow inhibition and strengthening fast inhibition will produce similar results to our own. Additionally, the LN-to-PN ratio could vary considerably within our model while yielding the same results as long as the synaptic strengths are adjusted accordingly. In a real honey bee, the AL has many more cells, so connectivity variability across neurons is much lower. The variability driving the results of this study, which uses the number of presynaptic LNs as a proxy for incoming slow inhibition strength onto PNs (Fig. 3), could instead arise from variation in synaptic weights between AL neurons. Thus, our parameter choices represent a single option out of a wide range of values that can be used to model experimentally observed AL behavior.

### 4.3 Stimulus modeling

Odor and wind stimuli were modeled as spiking Poisson processes. For PN *j* in the network, an incoming spike was modeled as an instantaneous jump of 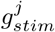 of magnitude 0.004 or 0.0033 if neuron *j* was an LN, followed by a decay time of *τ*_*stim*_ = 2 ms (for both LNs and PNs). We modeled three sources of stimuli: background noise at a constant rate of *λ*_*back*_ = 3.3 spikes/sec, odor input that had a maximum rate of 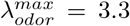 spikes/sec, and mechanosensory input at a rate of 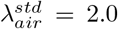 spikes/sec, unless otherwise noted.The total incoming input to cell *j* was given by:

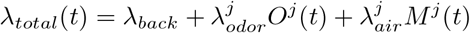

where *O*^*j*^(*t*) and *M*^*j*^(*t*) are functions that range between 0 and 1 used to model the temporal structure of the stimuli.

To simulate background noise alone, we set 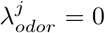 and 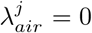 for all cells for the duration of the simulation. To simulate mechanosensory input alone in the time period [*t*_*on*_, *t*_*off*_], we set 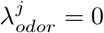 for all cells and 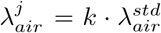 for all cells where *k* is positive real number (constant during the simulation) to denote different strengths of wind pulses. A single odor without mechanosensory input was modeled as an input pulse during the time period [*t*_*on*_, *t*_*off*_] to all cells in 3 out of 6 glomeruli (specific glomerular subset indicating odor identity). If neuron *j* was in a stimulated glomerulus, we set 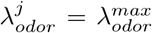; if neuron *j* was not in a stimulated glomerulus, we set 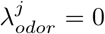. Since no mechanosensory input was present in this scenario, we set 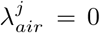 for all cells. Thus, mechanosensory input acts as a global stimulus, whereas odor input targets specific glomeruli. To simulate both mechanosensory and odor input, we employed a simple additive paradigm. If neuron *j* was in a glomeruli stimulated by an odor, 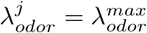 and we set 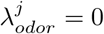 for all neurons not in this subset. Lastly, we set 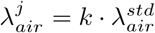 for all neurons in all glomeruli.

The function *M*^*j*^(*t*) represents the temporal dynamics of a mechanosensory stimulus pulse beginning at *t*_*on*_ and ending at *t*_*off*_ (in this work, *t*_*off*_ − *t*_*on*_ = 1000 ms). When *t < t*_*on*_ (where *t* is simulation time) *M*^*j*^(*t*) = 0; at *t* = *t*_*on*_, *M*^*j*^(*t*) increases from 0 to 1 sigmoidally if *j* is a PN, or instantly, if *j* is an LN. At time *t* = *t*_*off*_, *M*^*j*^(*t*) exponentially decays with a prescribed neuron dependent decay time. If neuron *j* was a PN, *M*^*j*^(*t*) had a rise time of *τ*_*rise*_ = 85 ms and a decay time of *τ*_*decay*_ = 400 ms:

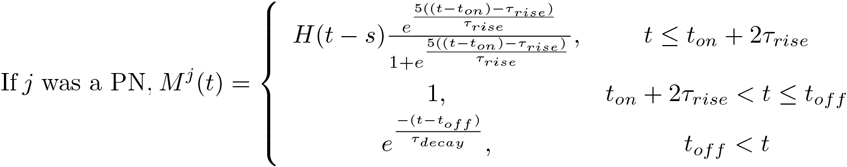

If neuron *j* was an LN, rise was instant while decay was still exponential with *τ*_*decay*_ = 150 ms:

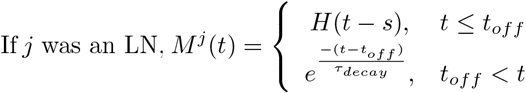

Similarly, *O*^*j*^(*t*) represents the temporal dynamics of odor stimulus. This was modeled in the same way. If neuron *j* was a PN, *O*^*j*^(*t*) had a rise time of *τ*_*rise*_ = 85 ms and a decay time of *τ*_*decay*_ = 450 ms:

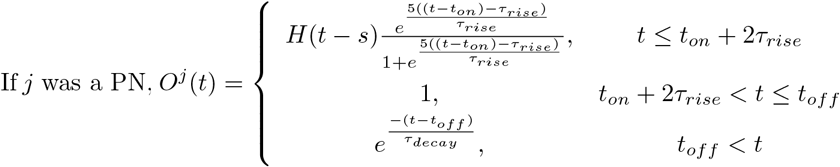

If neuron *j* was an LN, rise was instant while decay was still exponential with *τ*_*decay*_ = 200 ms:

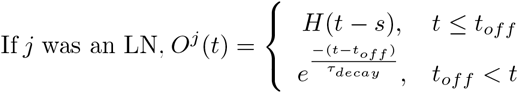

Again, the rise and decay time of both *O*^*j*^(*t*) and *M*^*j*^(*t*) were significantly longer if neuron *j* was a PN.

### 4.4 Data analysis

We constructed an algorithm to sort PNs into five response classes - none, sustained, transient, biphasic, and offset - based on trial-averaged firing rates in 50 ms bins. Since the stimulus rise and decay times were fixed parameters for all LNs and PNs, we could accurately predict the time intervals of PN stimulus-induced spiking. In our model, mechanosensory input was simulated as a 1 second pulse beginning at 400 ms and ending at 1400 ms. PN onset firing, denoted *v*_*on*_, was defined as the average firing rate in the interval [500 ms, 600 ms]; sustained firing, *v*_*sust*_, was the average firing rate in the interval [650 ms, 1350 ms]; offset firing, *v*_*off*_, was the average firing rate in the interval [1550 ms, 1650 ms]. Further, let *v*_*back*_ denote the average firing rate in the absence of stimulus, and let *σ*_*back*_, *σ*_*on*_, and *σ*_*sust*_ denote the standard deviations of PN firing rates during background, onset, and sustained periods, respectively. We heuristically categorized the response pattern of PN *j* by comparing these variables in this order:

1. Biphasic if 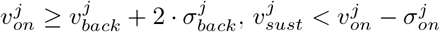, and 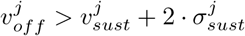.
2. Offset if 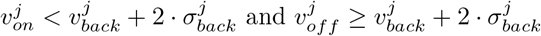.
3. None if 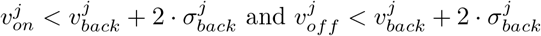.
4. Transient if 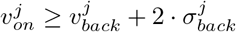 and 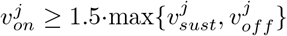.
5. Sustained if none of the above criteria are met.

In Fig. 8, we investigated the ability of our AL network to separate similar odors by comparing the distances between input and output vectors. The procedure was as follows. Two similar odors were simulated separately with and without mechanosensory stimulus: odor 1 to glomeruli 1-3, odor 2 to glomeruli 2-4. Input vectors were a 50 ms window average of the excitatory stimulus delivered to each PN in the network (60 total). Let 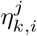 denote the average input to PN *j* at time interval *i* from odor *k* ∈ {1, 2}. Output vectors were the 50-trial averaged PN firing rates in 50 ms bins. Let 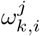 denote the average firing rate of PN *j* at time interval *i* under odor *k*. We then normalized each vector to one:

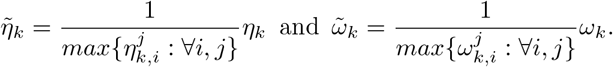

To find the distance between the input vectors of the two odors at time *i*, denoted 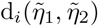, we used the Euclidean metric:

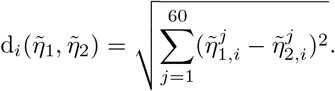

The distance between output vectors was calculated in the same way. This method was then repeated with mechanosensory input present.

Numerical integration was carried out using the Euler Method with a timestep of Δ*t* = 0.1 ms. All code, data analysis, and figures were written and plotted in MATLAB.

## Acknowledgements

Patel was supported by a grant from the National Institutes of Health (NIH R01DC020892). Reed was supported by a honors summer research fellowship from the Charles Center at William & Mary. We would like to thank Hong Lei and Shawn Mahoney at Arizona State University, our experimental collaborators on this project.

